# The evolutionary game of interspecific mutualism in the multi-species model

**DOI:** 10.1101/335133

**Authors:** Shota Shibasaki

**Affiliations:** Department of General Systems Studies, Graduate School of Arts and Sciences, The University of Tokyo, Tokyo, 1538902, Japan

**Keywords:** evolutionary game theory (EGT), mutualism, Red Queen effect, Red King effect

## Abstract

Mutualistic interspecific interactions, including Müllerian mimicry and division of labor, are common in nature. In contrast to antagonistic interactions, where faster evolution is favored, mutualism can favor slower evolution under some conditions. This is called the Red King effect. After Bergstrom and Lachmann (2003) proposed Red King effect, it has been investigated in two species models. However, biological examples suggest that the mutualisms can include three or more species. Here, I modeled the evolutionary dynamics of mutualism in communities where there exist two or more species, and all species mutually interact with one another. Regardless of the number of species in the community, it is possible to derive conditions for the stable equilibria. Although there exist nonlinear relationship between the evolutionary rates and the evolutionary fate of each species in the multi-species communities, the model suggests that it is possible to predict whether the faster evolution is favored or disfavored for the relatively fast species; on the other hand, it is difficult to predict the evolutionary fate of relatively slow species because the evolutionary dynamics of the slow species is affected by the evolutionary fate of the fast species.

## 1 Introduction

Mutualism, or cooperation between and among species, is widely spread in ecosystems. Two well known examples of mutualism are Müllerian mimicry and division of labor. In Müllerian mimicry, unpalatable species have evolved similar appearances, and they are each less likely to be predated upon because the predators effectively learn that these species are noxious (Muller, 1879). Although many empirical studies deal with Müllerian mimicry in butterflies (Sherratt, 2008), other examples include moths (Sbordoni et al., 1979; Niehuis et al., 2007), poison frogs (Chiari et al., 2004), vipers (Sanders et al., 2006), and fish (Springer and Smith-Vaniz, 1972). In division of labor, on the other hand, each species specializes certain tasks and exchanges different goods or services (Leigh, 2010). The examples of division of labor are found in the relationships within the gut microbiota (Rakoff-Nahoum et al., 2016), between plants and mycorrhizae (Remy et al., 1994), and between ants and aphids (Way, 1963).

Conflict can arise in mutualism regarding the role of each species. In Muullerian mimicry, it could be more advantageous to be a model species than a mimic because the life cycles, the habitats, and the body plans of model species are innate while a mimic species has to change these aspects to mimic (Veller et al., 2017). In division of labor, conflict can arise regarding species tasks (Wahl, 2002); when providing nutrients, one type of nutrient might have a greater cost for the organism than another type of nutrient. Such mutualistic symbioses with varying degrees of conflict have been conceptualized by the by-matrix snowdrift game whose payoff matrix is given by Table 1, where 0 ≤ *k* ≤ 2 because of the conflict between the species. In this game, each species performs either a generous strategy or a selfish strategy, and the left payoff in each cell is for species *i* and the right payoff is for species *j*. Nash equilibria are, therefore, where one species plays the generous strategy and the other species play the selfish strategy.

**Table 1:** Payoff matrix of the by-matrix snowdrift game

| | | species $j$ | |
| --- | --- | --- | --- |
|  |  | Generous | Selfish |
| species $i$ | Generous | $(k, k)$ | $(1, 2)$ |
| | Selfish | $(2, 1)$ | $(0, 0)$ |

Bergstrom and Lachmann (2003) modeled the evolutionary dynamics of the two species mutualism with a degree of conflict and they found that the slower evolving species is more likely to reach a favorable equilibrium than the faster species under some conditions. The slower evolutionary rate is caused, for example, by the longer generation time or the smaller mutation rate. The authors called this effect the Red King (RK) effect, which is converse to the Red Queen (RQ) effect (Van Valen, 1973), where the faster evolution is favored in antagonistic symbioses (and sometimes mutualistic symbioses as noted by Herre et al. (1999)). Although the model analyzed by Bergstrom and Lachmann was very simple (i.e., assuming a two-player game and infinite population sizes for both species), other researchers relaxed these assumptions and investigated the RK and/or RQ effects in mutualism with a degree of conflict. Gokhale and Traulsen (2012) found that in the multi-player snowdrift game of the two-species model, RK effect can change to the RQ effect. Gao et al. (2015) uncovered that a reward mechanism in the multi-payer snowdrift game causes the shift between the RK and RQ effects. Veller et al. (2017) investigated the factors which changes the evolutionary rates of each species (generation time, mutation rates, selection strength, and population sizes) in finite population, and they found that the RK effect shift to the RQ effect and vice versa from the short time scale to the long time scale due to the stochasticity.

Although these previous studies on the RK effect assume two-species communities, mutualisms can include more than two species. In the context of Müllerian mimicry, such phenomenon is called as the (Müllerian) mimicry ring, where three or more unpalatable species show the similar appearances to avoid predation (Sherratt, 2008). Examples of Müllerian mimicry rings include Appalachian millipedes (Marek and Bond, 2009), bumble bees (Plowright and Owen, 1980), cotton-stainer bugs (Zrzavý and Nedved, 1999), and *Heliconius* butterflies (Mallet and Gilbert, 1995). In the context of division of labor, as well, mutualistic symbioses are not limited to one-to-one relationships. For example, green algae can display mutualism with with several phylogenetically broad fungal species (Hom and Murray, 2014).

Inspired by these biological examples, I investigated whether it is possible to predict whether the faster evolution or slower evolution is favored in communities that include more than two species. Although the nonlinearity arises in the multi-species communities and it is difficult to say whether the faster evolution or the slower evolution is favored in the entire communities, the model suggests that it is predictable whether the faster or the slower evolution is favored for only the relatively fast species, especially if many species coexist in the communities.

## 2 Models

In this paper, I extend the original model of the Red King effect (Bergstrom and Lachmann, 2003) by generalizing the number of species in a community. Mutualistic symbioses with a degree of conflict are conceptualized by the by-matrix snowdrift game, whose payoff matrix is given by Table 1.

In this model, the fitness of each species is determined only by the interspecific interactions (i.e., intraspecific interactions are ignored). Given the number of species in the community *M*, the evolutionary dynamics of the fraction of generous individuals in species *i* is given by the replicator dynamics as below:
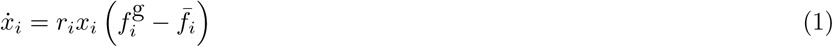

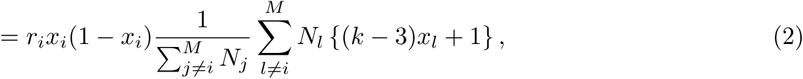

where *r_i_* is the evolutionary rate of species *i*, 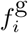 is the mean fitness of the generous individuals in species *i*, 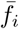 is the mean fitness of species *i*, and *N*. is the relative population size of species *i*. The parameters in this model are summarized in Table 2. In this model, the intraspecific interaction is ignored and the interspecific interactions are weighted by the population sizes. Notice that assumptions on the initial conditions, or the initial fractions of generous individuals in each species are important because the evolutionary dynamics is affected by the initial conditions.

**Table 2:** The parameters and variables in the model

| Notation | Interval | Description |
| --- | --- | --- |
| $x_i$ | $\in [0, 1]$ | Fraction of generous individuals in species $i$ |
| $r_i$ | $> 0$ | Evolutionary rate of species $i$ |
| $N_i$ | $> 0$ | Relative population size of species $i$ |
| $M$ | $\geq 2$ (integer) | Number of species in the community |

For clarity, I explicitly define Red King and Red Queen effects here. For convenience, I call species *i* generous (selfish) species when all individuals of species *i* become generous (selfish) at equilibrium states: i.e., the probabilities of 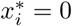, and 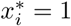, respectively.

**Definitions of the Red King and Red Queen effects**. *Given that the initial fractions of the generous individuals in each species x_i_*(0) *are uniformly independent and identically distributed (i.i.d.), one can say the Red King (Red Queen) effect is operating when the slower (the faster) the species evolves, the more likely it is to become a selfish species. In other words, if the RK or RQ effect operates, there should be a linear relationship between the order of the evolutionary rates and the probability that each species become selfish*.

Although previous studies on the RK effect (Bergstrom and Lachmann, 2003; Gokhale and Traulsen, 2012) measured the sizes of basin of attraction wherein either of the two species becomes selfish, it is difficult to analyze in the same way in *M* species communities due to the high dimensionality of the model and the existence of multiple stable equilibria (see Results). To avoid this problem, the favorabilities were measured by the probabilities that focal species evolved selfishly in this study. In addition, it is necessary to consider the probability density of the initial conditions (*x*_1_(0),…, *x_M_*(0)) because the initial conditions determine which equilibrium states the communities converge to. In other words, the probability density of the initial conditions affects the favorabilites of the species in the communities although the evolutionary dynamics given by Eq (2) is deterministic. In this model, I assumed that the initial fractions of the generous individuals are uniformly *i*.i.d. and no particular initial condition is weighted, but other assumptions on the initial conditions will change the results.

## 3 Results

In this section, I shall show the conditions for linear stable equilibria and analyze the effect of the evolutionary rates on the evolutionary fate of each species. First, the conditions for the stable equilibrium state is derived, and then, the relationship between the evolutionary rates and the favorabilities using the computer simulations. In the communities with a large value of *M*, however, the computational cost of the simulation is large. To avoid this problem, the analytical results under the ideal conditions are shown.

### 3.1 Stable equilibria

Assuming that the population size of each species is infinitely large (*N_i_* → ∞ for *i* = 1,…, *M*), the evolutionary dynamics is represented as

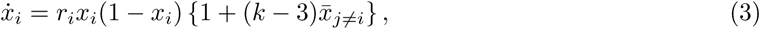

where 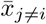 is the average fraction of generous individuals except for species *i*:
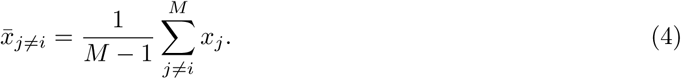

Notice that equation (3) is the same as the evolutionary dynamics proposed by Bergstrom and Lachmann (2003) when *M* = 2. The situation where the assumption on the population size is relaxed is analyzed in Appendix B.

In the stable equilibria in equation (3), there exist *m* generous species and *M* − *m* selfish species. The number of generous species *m* should satisfy the inequality below:
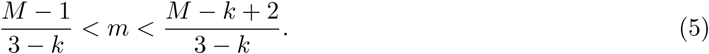

The derivation is shown in Appendix A. It should be noted that there exists at most one integer *m* that satisfies inequality (5) given the values of *k* and *M*. In a community with three species (*M* = 3), for example, one species becomes generous (*m* = 1) and other two species are selfish at stable equilibria if *k* is small (0 ≤ *k* ≤ 1); on the other hand, there exist two generous species (*m* = 2) and one selfish species at a stable equilibrium if *k* is large (1 < *k* < 2).

### 3.2 Computer simulation

Although inequality (5) indicates that there exist 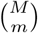 stable equilibria in the dynamics defined by equation (3), it is unclear whether species *i* is more likely to evolve generously or selfishly because the initial conditions determine which stable equilibria the dynamics converge to. To clarify this point, the evolutionary dynamics in the three and four species communities were simulated from various initial conditions. The evolutionary rate of each species is given by ***r*** = (1/8, 1, 8) in the three species model, and ***r*** = (1/8, 1/2, 2, 8) in the four species model, respectively. The initial conditions are given by changing the fraction of generous individuals in each species (the step size 0.05). The total numbers of simulations in the three and four species models are 8, 000 and 16, 000, respectively. The favorabilities, or the probabilities that each species evolve selfishly at the stable equilibrium is shown in Fig. 1.

**Figure 1:**
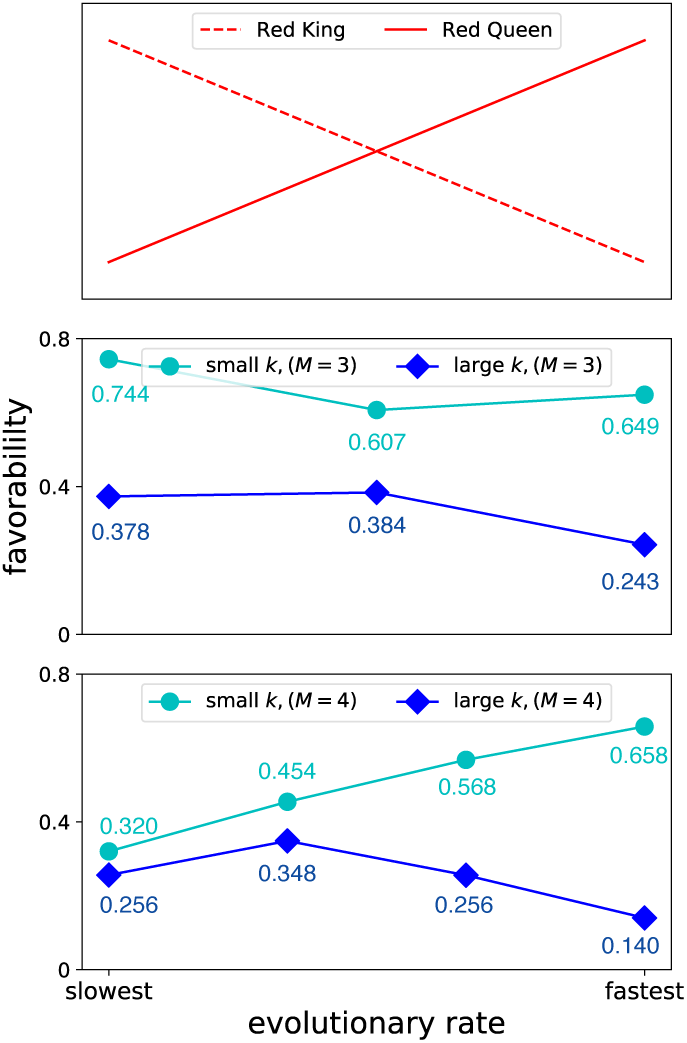
The evolutionary rates and the probability of becoming selfish species. The relationship between the order of the evolutionary rates and the probability that each species becomes selfish at stable equilibria (i.e., favorability) are shown. Top: the hypothetical results if RK effect or RQ effect operates; there should be a negative (positive) relationships between the order of evolutionary rates and the favorability. However, the results of computer simulations (Middle: three species, and Bottom: four species) did not show such relationships except for the case of small *k* in the four species community. The parameters are: ***r*** = (1/8, 1, 8), and *k* = 0.5 (small) or *k* = 1.5 (large) in the three species model, and ***r*** = (1/8, 1/2, 2, 8) and *k* = 0.5 (small) or *k* = 1.6 (large) in the four species model.

Except for the case of the four species model with small *k*, there is no clear RQ effect nor RK effect in Fig. 1; there exists nonlinear relationship between the evolutionary rates and probabilities that each species becomes selfish (favorability); in the three species community, the slowly evolving species and the fast species are more favorable than the intermediate species if *k* is small. When *k* is large in the three species model, there exist no clear RQ nor RK effects either, although the difference of the probabilities of being selfish is small between the slowly evolving species and the intermediate one. In the four species model, there exist a convex relationship between the order of the evolutionary rates and the favorability when *k* is large. On the other hand, there exists a positive relationship and, therefore, a RQ effect if *k* is small. These results suggest there can be the mixture of the RQ effect and the RK effect in multi-species communities.

The analysis with the computer simulation is, however, ineffective when the number of species in the community *M* is large; although the numerical integration of the evolutionary dynamics given by Eq (3) does not take a long time, the space of the initial conditions enlarge as *M* increases. In other words, the total computational cost increases when *M* increases. One way to avoid this problem is assuming an ideal condition where only one species changes the fraction of generous individuals until this species fixes its strategy while the other species do not evolve. Under this ideal condition, the value of *Xj=_i_* does not change until species *i* fixes it’s strategy, and therefore, the evolutionary fate of species *i* is determined by the sign of 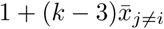 (positive: generous, and negative: selfish).

The computer simulations suggested that the order of fixation is consist with the order of the evolutionary rate (Fig. 2); for example, the fastest species is the species that is the most likely to the strategy at first. In other words, the effect of the initial conditions on the order of the fixation isnegligibly small, and the the order of the fixation is predictable from the order of the evolutionary rate.

**Figure 2:**
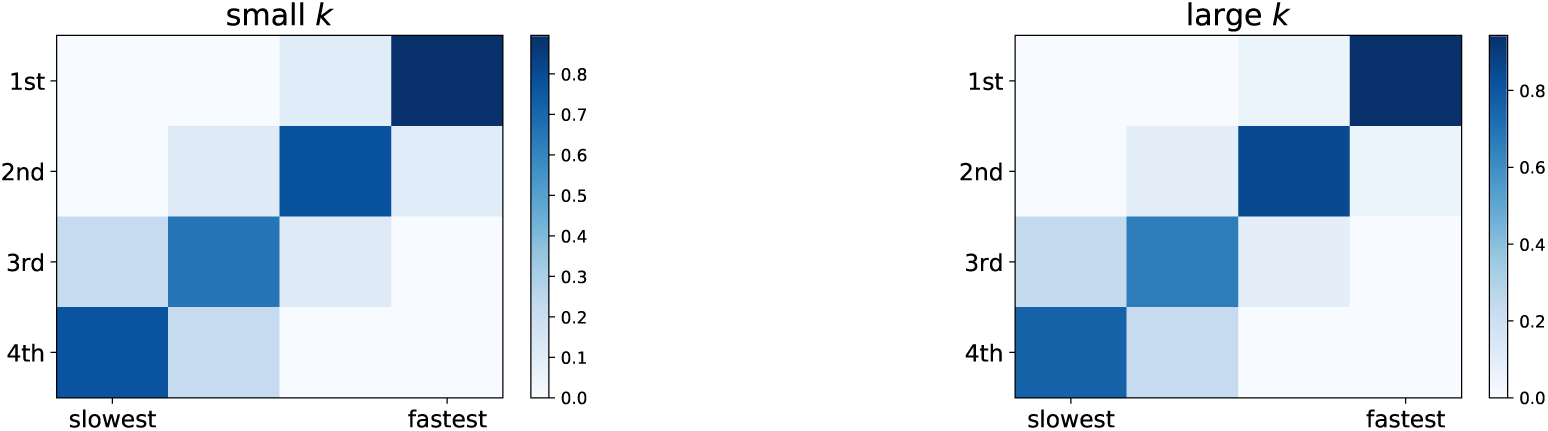
The order of the evolutionary rates and the order of the fixation. The relationship between the evolutionary rates (horizontal axis) of each species and the order of the fixation (vertical axis) in the four species model with a small value of *k* (*k* = 0.5, left) and a large value (*k* = 1.6, right). The color of each cell represent the probability that focal species fixes its strategy at the given order. Regardless of the value of *k*,there exist consistency between the order of the evolutionary rates and the order of the fixation; the species that is the most likely to fix its strategy at first is the fastest species, and those that fixes the strategy at last is the species with the slowest evolutionary rate. The evolutionary rates and the initial conditions are the same as in Fig. 1.

### 3.3 Analysis under the ideal condition

As the order of the fixation can be estimated by the order of the evolutionary rate, let us assume the ideal condition where the differences in evolutionary rates between each species are quite large. Under this condition, only the fastest evolving species that has never fixed its strategy can evolve toward generosity or selfishness while the remaining species do not change in their fractions of generous individuals. In other words, each species fixes its strategy according to the order of the evolutionary rate, and the evolutionary direction of each species is determined by the initial condition and the evolutionary fate of the species which have already fixed their strategies because the sign of 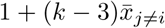 determines the evolutionary direction. Notice that the it is necessary to the calculate the probability distribution of the initial conditions because the evolutionary dynamics is affected by the initial conditions, although the dynamics is deterministic.

When species *i* is the *i*th fastest species in the *M* species community (*i* = 1,…, *M* − 1), the focal species *i* is more likely to evolve generously or selfishly if and only if

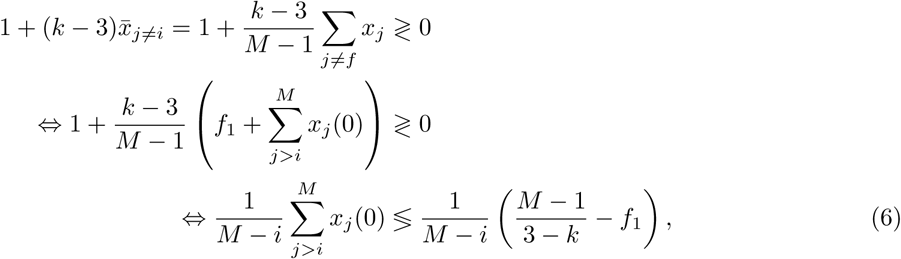

where *f*_1_ is the number of species which have already been fixed as generous species. Notice that it is not necessary to consider the slowest species *i* = *M* because the evolutionary fate of the slowest species is determined only by the evolutionary fate of the other species (if there already exist *m* generous species, the slowest species become selfish; otherwise, the slowest species evolve generously). The upper (lower) sign of Eq (6) is the case when species *i* is more likely to be generous (selfish) species. As the initial fraction of generous individuals in each species is uniformly i.i.d., the left-hand side of Eq (6) represents the mean of *M* − *i* independent samples from the uniform distribution. Using the central limit theorem, the left-hand side of Eq(6) approximately follows the normal distribution 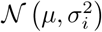 whose mean is *μ* = 0, 5 and the inverse of the variance is 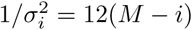.

The right-hand side of Eq (6), on the other hand, can be regarded as the threshold for species *i T_i_*; if the left-hand side of Eq (6) is smaller (larger) than *T_i_*, species *i* evolves generously (selfishly). This threshold is, however, affected by the evolutionary fates of the faster species, which have already fixed their strategies. By denoting the number of species which have already been fixed as selfish *f*_1_ = *i* + 1 − *f*_0_, the right-hand side of Eq (6) is seen as the threshold for the evolutionary direction of species *i*, whose value is determined by *f*_0_, *f*_1_;
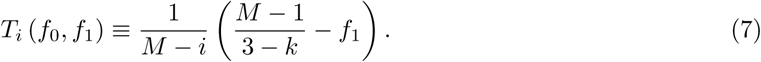

From Eqs(6) and (7), the favorability of species *i* is computable with arbitrary values of *k* and *M*. Given the value of *f*_0_ and *f*_1_, let denote the conditional probability that species *i* become selfish as 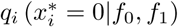. Then, the favorability of species *i* is written as

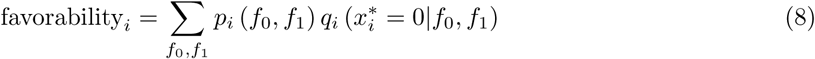

where *p_i_* (*f*_0_, *f*_1_) is the probability density of (*f*_0_, *f*_1_) for species *i*. From Eq (6), the conditional probability 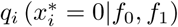 is derived as

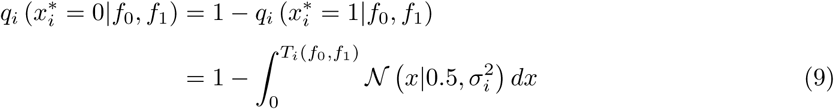

where 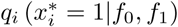 indicates the conditional probability that species *i* become generous given the value of (*f*_0_, *f*_1_). Notice that these two conditional probabilities are affected by the values of *f*_0_ and *f*_1_, but given the values of *f*_0_ and *f*_1_, the two conditional probabilities can easily be computed by calculating the cumulative probability function of the normal distribution 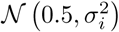.

The probability density *p_i_* (*f*_0_, *f*_1_) is, on the other hand, sequentially computable (Fig. 3). For the fastest species (*i* = 1), *p*_1_ (0, 0) = 1 and the favorability of the fastest species is derived by

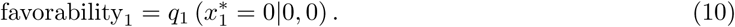

To calculate the favorability of the second fastest species, on the other hand, it is necessary to calculate the probability densities of (*f*_0_, *f*_1_) = (0, 1), (1, 0). However, the probability that (*f*_0_, *f*_1_) = (0,1) (or (*f*_0_, *f*_1_) = (1, 0)) is the same as the probability that the fastest species become generous (selfish), respectively. The favorability of the second fastest species is, therefore, derived as below:
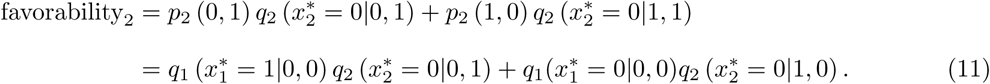

For the third fastest species (*i* = 3), the probability density of (*f*_0_, *f*_1_) is derived as follows:
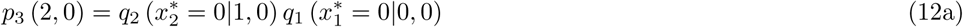

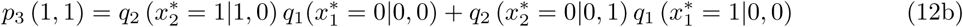

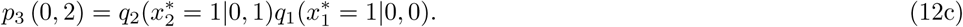

By substituting Eq (7) into Eq (9), the conditional probabilities that the third fastest becomes selfish species are achieved. Then, the favorability of the third fastest species is derived from Eq (8). Therefore, for any value of *i*, the favorability of species *i* is computable by the conditional probabilities that the species evolving faster than *i* becomes selfish.

Fig. 4 shows the results of analysis on the relationship between the evolutionary rates and the favorabilities under the ideal conditions. As the results of the computer simulations (Fig. 1), there exist nonlinear relationships between the evolutionary rates and the favorabilities. Moreover, in the four species model with the small value of *k*, where the computer simulation represents the linear relationships between the evolutionary rates and the favorabilities, the analysis under the ideal conditions shows the nonlinear relationship.

**Figure 3:**
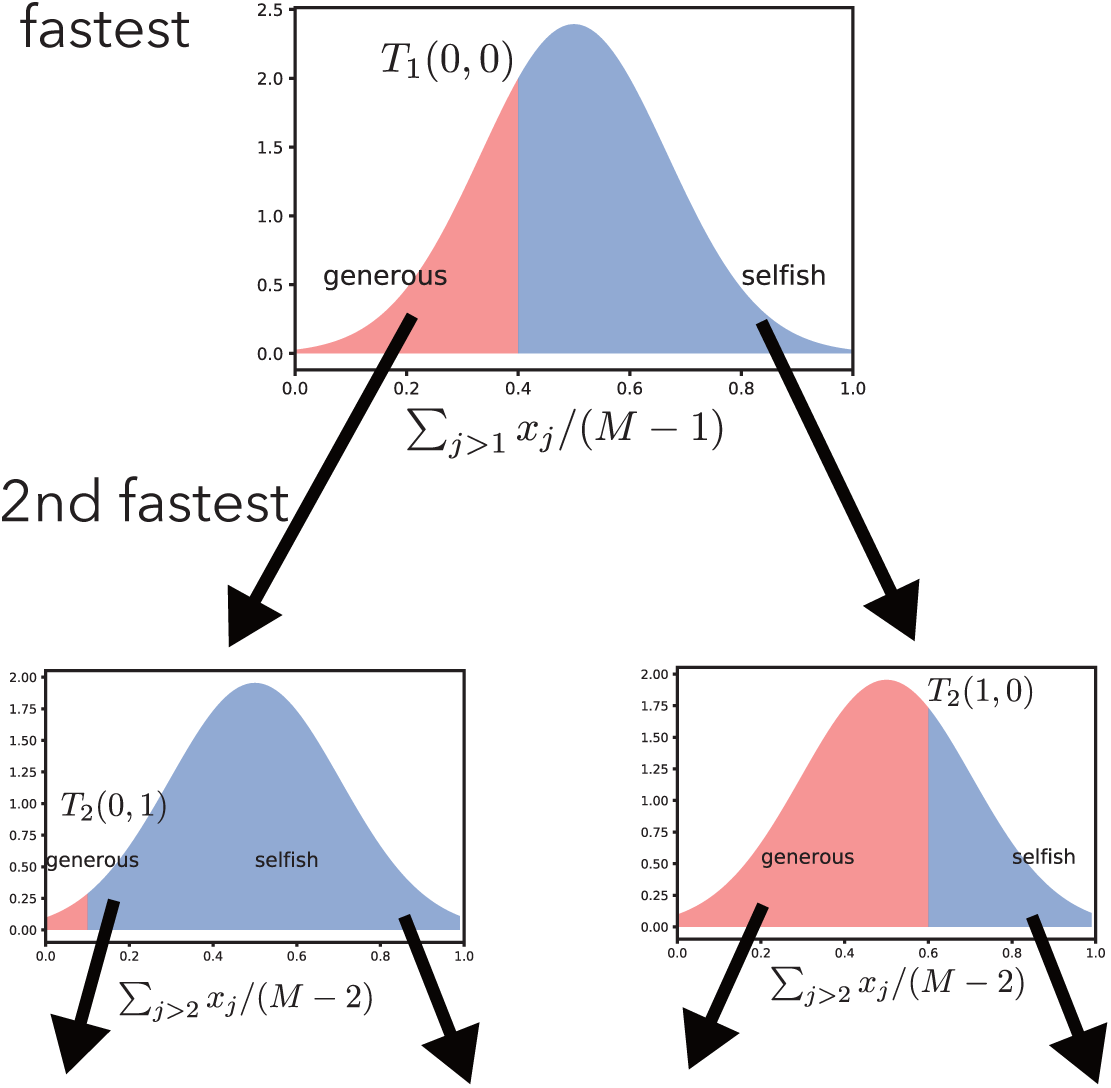
The schematic illustration of the analysis under the ideal conditions. The threshold of the evolutionary direction is determined by the evolutionary fate of the species which have already fixed their strategies. Under the ideal conditions, the evolutionary fate of the fastest species (*i* = 1) is determined only by the value of *k*. In other words, the favorability of the fastest species is computable with arbitrary value of and *k*. The threshold for the second fastest species (*i* = 2) is determined not only by *k* but also the strategy of the fastest species (left: generous and right: selfish). Notice that the probability *p*_2_ (0, 1) (*p*_2_ (1, 0))is the same as the probability that the fastest species becomes generous (selfish). The favorability of the second fastest species is, therefore, also computable. In addition, from the computation above, the probability density of (*f*_0_, *f*_1_) for the third fastest species *p*_3_ (*f*_0_, *f*_1_) is calculated.

The advantage of the analysis under the ideal conditions is that the ideal conditions enable to analyze the communities with larger number of species., which requires the quite large computational costs in the computer simulation due to the high dimensionality. Although there still exist nonlinear relationships between the evolutionary rates and the favorabilities when *M* is large (e.g., *M* = 20), there exist a pattern which cannot be seen when *M* is small (Fig. 5); if the focal species has a relatively fast evolutionary rate, the faster the species evolves, the larger (the smaller) the favorability of the species is when *k* is small (large). The species with relatively intermediate evolutionary rates show the almost same favorabilities (around 0.5), which means that the difference of the evolutionary rates has little effect on the favorabilities for these species. The species with the relatively slowly evolving species, especially the slowest species, however, it is difficult to find a pattern. To sum up, under the ideal conditions, the communities with large number of *M* can show the pattern between the evolutionary rates and the favorabilities of the species with the relatively fast or intermediate evolutionary rates.

**Figure 4:**
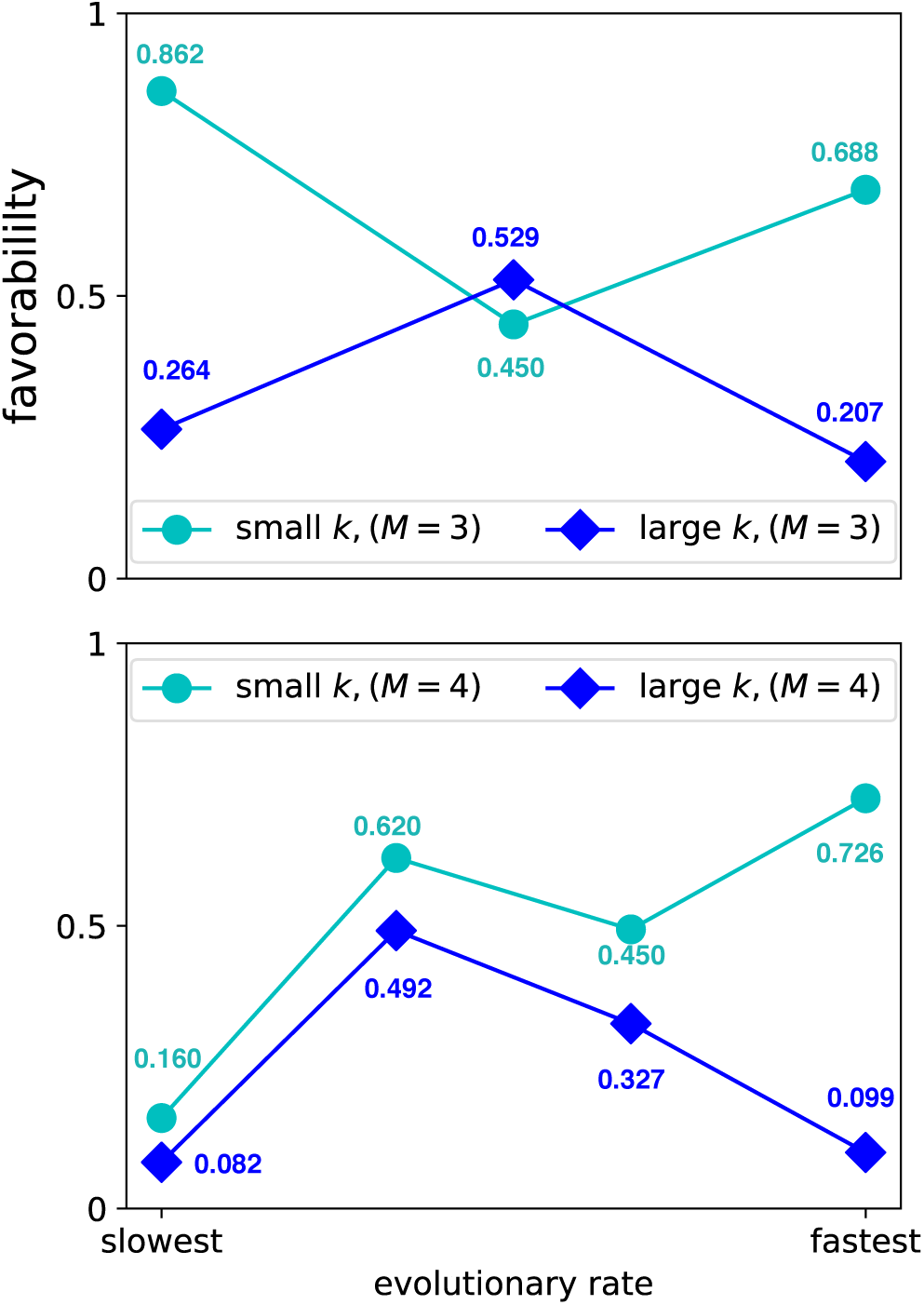
Favorabilities under the ideal conditions. The favorabilities of each species in the three species communities (*M* = 3) and the four species communities (*M* = 4) are computed under the ideal conditions. using different value of *k* (in the three species communities, *k* = 0.5 for small *k* and *k* = 1.5 for large *k* whereas *k* = 0.5 for small *k* and *k* = 1.6 for large *k* in the four species community). It is assumed that the left-hand side of Inequality (6) is distributed according to the normal distribution if *i* = 1, 2,…,*M* − 2; otherwise the uniform distribution is used for the distribution of the left hand side in inequality (6).

## 4 Discussion

In this paper, the effect of evolutionary rates on mutualism is investigated by generalizing the model proposed by (Bergstrom and Lachmann, 2003). In particular, I modified their model with respect to of the number of species in the communities and the population sizes of each species. Although the evolutionary dynamics is deterministic, the stable equilibrium where the dynamics converges depends on the initial conditions, and therefore, the favorability or the probability that the focal species become selfish at a stable equilibrium should be evaluated.

**Figure 5:**
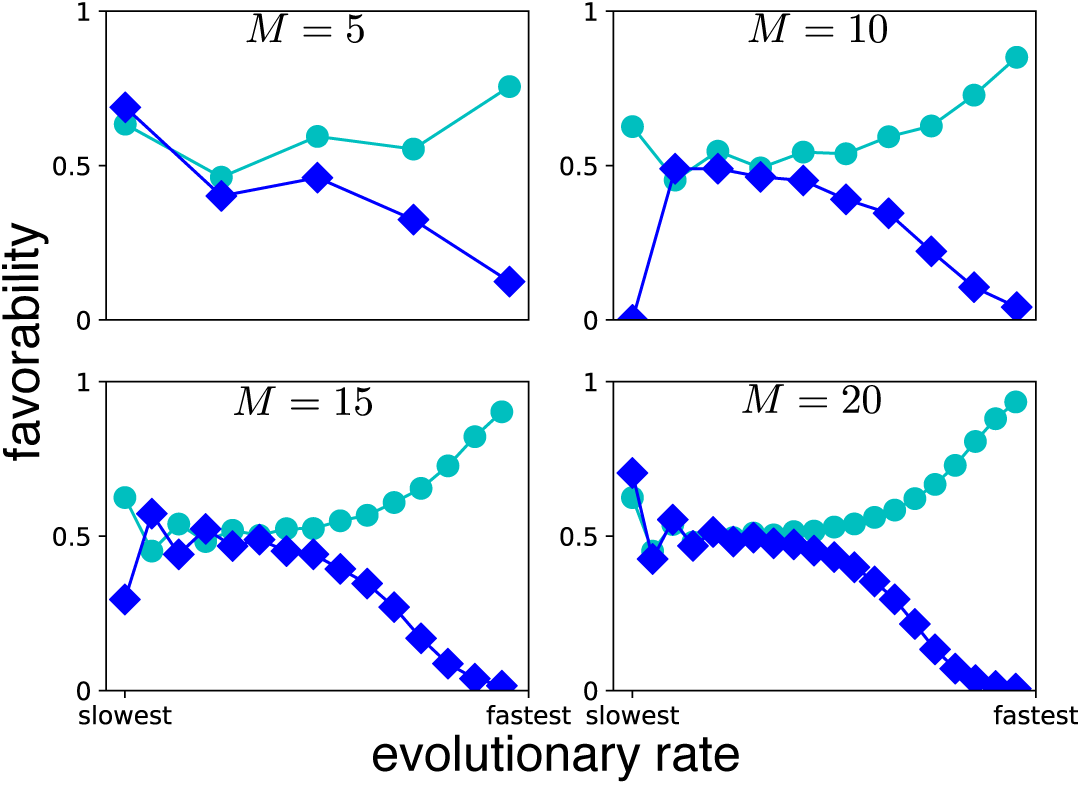
Favorabilities under the ideal conditions in larger communities. The favorabilities under the ideal conditions when *M* = 5,10,15, and 20 are shown. In each value of *M*, the value of *k* is changed (cyan: *k* = 0.5, and blue *k* = 1.5). Although there exist nonlinear relationships between the evolutionary rates and the favorabilities, the models with large value of *M* show that the faster the evolutionary rate is, the more the focal species is likely to evolve generously (selfishly) if *k* = 0.5 (*k* =1.5) and if the focal species has the enough large evolutionary rate.

In the original two-species model of Red King (Bergstrom and Lachmann, 2003), the stable equilibria are where one species become generous and the other species become selfish. At such stable equilibria, the selfish species receives a larger benefit than the generous species, and one may say the selfish species is a “winner” and the generous species is a “loser”. In the multi-species model of this paper, on the other hand, the selfish species are not always such “winners”. For example, in the three species model where all species have the same population size, there exist 1 generous and 2 selfish species at the stable equilibria when 0 ≤ *k <* 1, while the stable equilibria have 2 generous and 1 selfish species when 1 *< k <* 2. In the latter case, the selfish species receive 4 payoff while the generous species receive *k* + 1 payoff in each, meaning that the selfish species “wins” the both of the two generous species. However, if 0 ≤ *k* < 1, the payoffs of all species are 2; both the selfish species and the generous species receive the same amount of benefit; therefore, the two selfish species do not “win” the generous species.

Better interpretations of the generous strategy and the selfish one would be as follows; the generous strategy is a role which produces a benefit to other species (e.g., producing a leaky resource which works as energy for the producer species andother species), and the selfish strategy is a role which pays a small cost but does not produce benefits unless other species play the generous strategy (e.g., producing enzyme which makes the resource available more effectively). While the selfish species always receive a larger benefit from the generous species than the selfish one, the generous species receives more (or less) benefits from the selfish species than the generous one when 0 ≤ *k* < 1 (or < 1k < 2). In the context of the division of labor of producing the leaky resource and the enzyme, the diffusion rate of the enzyme can be a measure of *k*. For example, 0 ≤ *k* < 1 represents the case where the diffusion rate of the enzyme is large and both the generous species and the selfish species use the leaky resource effectively due tot the enzyme the selfish species produces. The difference in the payoffs of the two species would represent the differences in the cost of producing the resource or the enzyme. The situation where the diffusion rate of the enzyme is low, on the other hand, will be represented by the parameter 1 < *k* < 2. In such case, only the selfish species make the resource available effectively due to the benefit of the enzyme while the generous species cannot access the enzyme. In such situation, it is better for the generous species that other species play the generous strategy and increase the amount of the leaky resource. Considering this interpretation, the generous species can be regarded as one which provides the benefit of the mutualism through the community, while the selfish species can be regarded as one that selfishly maximizes its potential payoff.

While the two-species model of the two player game shows the positive or negative correlations between the evolutionary rates and the favorabilities (Bergstrom and Lachmann, 2003; Veller et al., 2017), the computer simulation (Fig. 1) and the analysis under the ideal conditions (Fig. 4) in this paper represent the nonlinear relationship between the evolutionary rates and the favorabilities in the three or four species communities. Although the computer simulations of the communities with the large value of *M* take a long time, the assumption of the ideal conditions enable to analyze the communities where many species coexist. Although the nonlinear relationship between the evolutionary rates and the favorabilities remains when *M* increases, one can find a clear pattern in the relationship (Fig. 5); the relatively fast species have the larger (smaller) favorabilities as the focal species evolves faster when *k* is small (*k* is large), while the favorabilities of the species with the relatively intermediate evolutionary rates are around 0.5. For the relatively slow species, however, it is difficult find any tendency.

Such pattern suggests that the faster evolution is favored (disfavored) for the relatively fast species in the community with many species if *k* is smaller (larger) than 1. This result is consistent with the original two-species model (Bergstrom and Lachmann, 2003), where the Red Queen effect (the King effect) operates when *k* < 1 (*k* > 1). This consistency would arise from the fact that faster evolving species is more sensitive to the value of *k*. Eq (7) suggests that the evolutionary direction is more sensitive to the values of *k*, which determines the evolutionary direction, when the focal species evolves faster (smaller *i*), because *f*_1_ is equal or smaller than *i* − 1. In other words, the evolutionary fate of the fast species is sensitive to the value of *k*, while the slow species is more sensitive to the evolutionary fate of the faster species, which would lead the difficulty of finding a pattern for the relatively slow species. Notice that an exception is a case when *M* = 2; if the evolutionary fate of the faster species is determined, then that of the slower species is also determined as there exist only stable equilibrium states. Therefore, in the multi-species communities, only the evolutionary fate of the relatively fast species is predictable from the value of *k*, the payoff when the generous species interacts with other generous species.

Of course, this research has some limitations. First, it would be unnatural that all species play the same game defined by Table 1. The parameter *k* could be replaced with *k_ij_*, which a generous individual of species *i* receives when interacting with a generous individual of species *j*. In addition, it is more natural if a community includes not only mutualistic interactions but also antagonistic interactions. Second, this paper assumes the two distinct strategies, selfish or generous. However, it is possible to consider continuous trait values where the payoff is then given by the difference of the trait values of the payers. Although the distinct strategies could be enough in the context of division of labor, continuous traits model is better in the context of mimicry as each species would have the different appearance and they have to pay different amount of cost to mimic the model species. Third, population sizes can change over time although the population sizes are fixed in this paper. The population dynamics can change the results in this study because the population size affects the evolutionary rates (Veller et al., 2017) and the stability of the equilibria, as shown in Appendix B. Indeed, recent studies combine the public good game with population dynamics and show the maintenance of cooperation (Hauert et al., 2006) and complex dynamics (Gokhale and Hauert, 2016). Such eco-evolutionary dynamics can also be analyzed in the mutualism with a degree of conflict.

In summary, this study analyzed the evolution of mutualism in the multi-species communities by generalizing the two-species model proposed by Bergstrom and Lachmann (2003). Although in the multi-species communities, there exist nonlinear relationship between the evolutionary rates and the favorabilities, it is possible to predict the evolutionary fates of relatively fast evolving species from the value of *k*, or the payoff of generous species when they interact with other generous individuals.

## Acknowledgement

The author thanks to Dr. P. Zee for his comments and English corrections in the earlier manuscript. The author also thanks Y. Otake for her beneficial comments to the earlier manuscript. The author is a Research Fellow of the Japan Society for the Promotion of Science (JSPS), and this work was supported by Grant-in-Aid for JSPS Fellows (grant number JP18J22388).

## Appendix A Stability analysis of infinite population sizes

Here, the linear stability of the equilibria in the model with infinite population sizes given by equation (3) is performed and I prove that stable equilibria are only those which has *m* generous species and *M* − *m* selfish species, where integer *m* satisfies inequality (5).

The element of the Jacobian matrix of equation (3) at the equilibrium ***x***^*^ is represented by

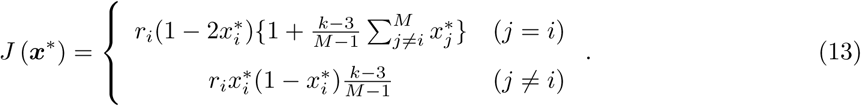

To analyze the stability of each equilibrium, I classified the equilibria into the two class: (i) an exterior equilibrium where all elements are either 0 or 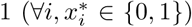, and (ii) an interior equilibrium which holds at least one element between 0 and 1 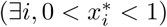.

## Appendix A.1 Stability of exterior equilibria

In the case of an exterior equilibrium 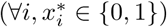, the Jacobian matrix *J* of such equilibrium is given by

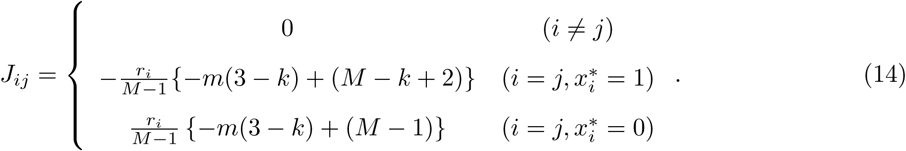

where *m* is the number of generous species at this equilibrium. As this Jacobian matrix is a diagonal matrix with an exterior equilibrium, the eigenvalues of the Jacobian matrix are the same as the diagonal elements of the Jacobian matrix. The exterior equilibrium is, therefore, linearly stable if and only if

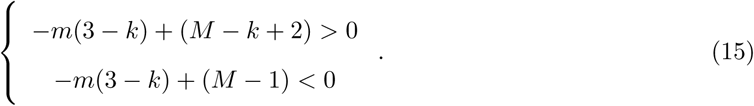

Here, inequalities (5) are obtained.

## Appendix A.2 Stability of interior equilibria

Next, let consider the stability of the interior equilibria 0 and 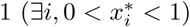. The interior equilibria can divide into two sub-categories: a full interior equilibrium where all elements are between 0 and 1, and a partially interior equilibrium where some elements are between 0 and 1 while the other elements are either 0 or 1. In this section, the full interior equilibrium is first analyzed because the stability analysis for the full interior equilibrium is simple. Then, the analysis for the partially interior equilibrium is shown.

There exist a unique full interior equilibrium in the evolutionary dynamics given by equation (3): ***x***^*^ = ((3 − *k*)^−1^; (3 − *k*)^−1^;…, (3 − *k*)^−1^. The Jacobian matrix at this equilibrium is written as

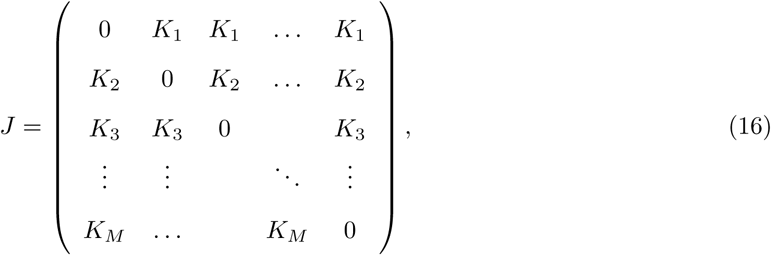

where *K_i_* = −*r_i_*(2 − *k*)/ {(3 − *k*)(*M* − 1)} ≤ 0. As the Jacobian matrix is not a diagonal matrix in this case, it is difficult to directly calculate the eigenvalues.

Instead of this approach, I shall show that Routh-Hurwitz criteria do not hold in the case of the full interior equilibrium and, therefore, the full interior equilibrium is not stable. Here, Mathematical Induction shows that the coefficient of *M* − 1th order in the characteristic equation is 0, which means that Routh-Hurwitz criteria do not hold (Murray, 2002, Appendix B.1).

### Proof.

First, let consider the case when *M* = 2. Then, the characteristic equation is written as

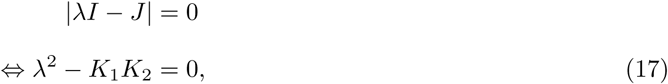

where *λ* is the eigenvalue of the Jacobian matrix given by equation (16), *I* is the identity matrix, and |*λI* − *J*| is the determinant of matrix *λI* − *J*. Obviously, the coefficient of λ is zero.

Next, let assume that the coefficient of *M* − 1th order in the characteristic equation is zero when *M* = 2; 3;…, *n*. If *M* = *n* + 1, the characteristic equation is

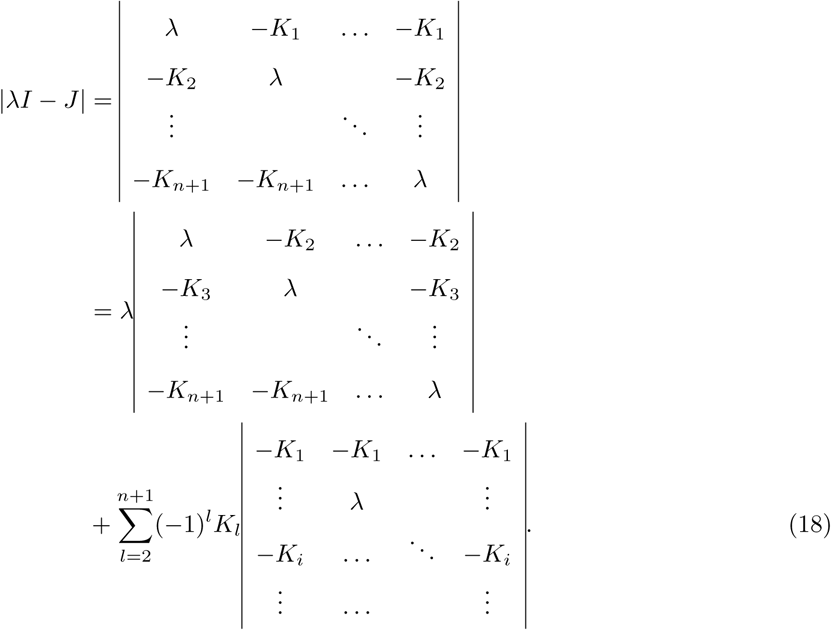

The first term in equation (18) is

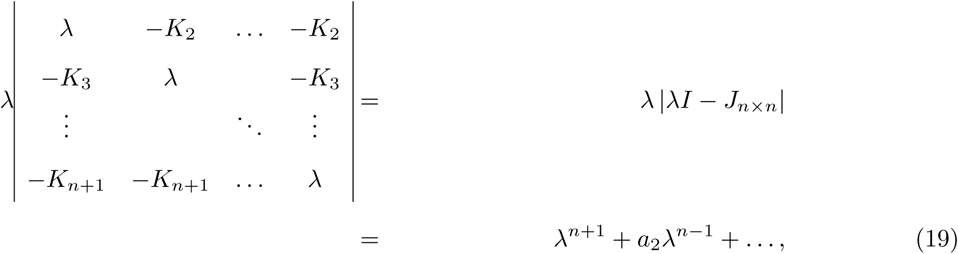

where *J_n_*_×_*_n_* is *n* × *n* matrix while *J* in equation (18) is (*n* +1) × (*n* + 1) matrix. The second term in equation (18) is, on the other hand, the *n* − 1th degree equation of *λ*. The coefficient of *n*th order in equation (18) is, therefor, zero.

From these result, Routh-Hurwitz criteria do not hold in the characteristic equation of the Jacobian matrix in equation (16) with the arbitrary integer of *M*. As all eigenvalues are negative if and only if the Routh-Hurwitz criteria hold, the full interior equilibrium is not stable.

Next, let consider the stability of the partially interior equilibria. At a partially interior equilibrium, I denoted the number of selfish and generous species as *f*_0_ and *f*_1_, respectively. The number of remaining species is *R* = *M* − (*f*_0_ + *f*_1_) and both generous individuals and selfish ones coexist within each species.

The elements of partially interior equilibrium is represented as below:
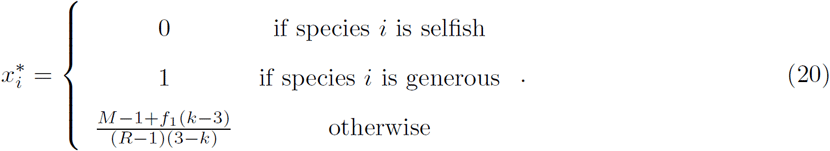

The Jacobian matrix of such equilibrium is written as equation (16), although *K_i_* = 0 if 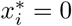 or 1. As the Mathematical Induction above again holds in this case, all partially interior equilibrium is not stable.

## Appendix B Stability analysis in the of different population size model

## Appendix B.1 Model formulation

In the main text, the analysis is based on the assumption that all species have the infinite population sizes. Here this assumption is relaxed and it is assumed that each species has a different population size. Notice that, however, the population sizes are enough large to use the ordinary differential equations and that the population sizes are not changed over time.

In the case that each species has a different population size, the evolutionary dynamics defined in equation (2) are written as

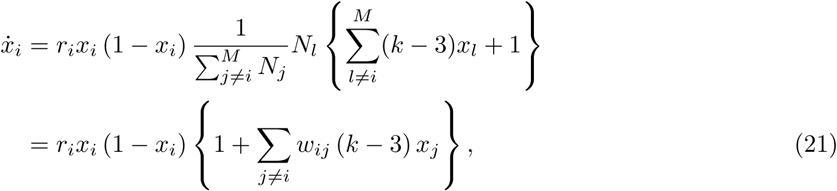

where 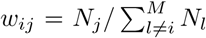. By giving *w_ii_* = 0 for *i* = 1, 2,…, *M*, one can find 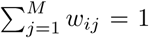 for any *i*. This means that *w_ij_*· is the weight of the effect of species *j* on species *i*. In this model, the results shown in the main text are rewritten by evaluating the weighted average of generous individuals 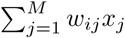.

The stable equilibria 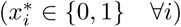 should satisfy the conditions below:
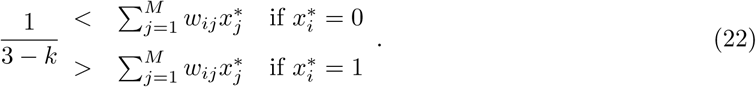

The derivation is shown in Appendix B.2. It should be noted that the population sizes do not affect the stability of each equilibrium in the case of *M* = 2. Inequalities (22) suggest that we need to evaluate matrix *W* = {*w_ij_*} and the value of *k* for all possible 2*^M^* combinations of the evolutionary fate of all species in the community. Interestingly, this model can hold the multiple stable equilibria which have different number of generous spices in each while the model where all species have infinite large population size shows that each stable equilibrium should have *m* generous species given the values of *k* and *M*. For example, there exist two stable equilibria and one has two generous and one selfish species while the other stable equilibrium has one generous and two selfish species when ***N*** = (1, 1, 5) and *k* = 0.5 (see Fig. 7d).

## Appendix B.2 Stability analysis

Here, I prove that in the model of different population sizes, the stable equilibria should satisfy inequalities (22) while the interior equilibria, which have at least one species wherein there exist both generous and selfish individuals cannot be stable. The elements of Jacobian matrix for equation (21) are represented as

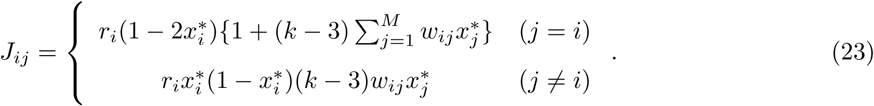

In the case of the exterior equilibria where all species have either generous or selfish individuals, Jacobian matrix is a diagonal matrix. An exterior equilibrium is stable if and only if all diagonal elements of Jacobian matrix are negative, which is the same as inequalities (22).

In the case of interior equilibria, on the other hand, all diagonal elements of Jacobian matrix are zeros. As shown in Appendix A.2, Routh-Hurwitz criteria do not hold in the characteristic equation with such Jacobian matrix, although the row elements except for the diagonal elements in each row do not have the same values in this case.

## Appendix C Code availability

The codes used in this research are available at Github.

## Appendix D Supplementary figures

**Figure 6:**
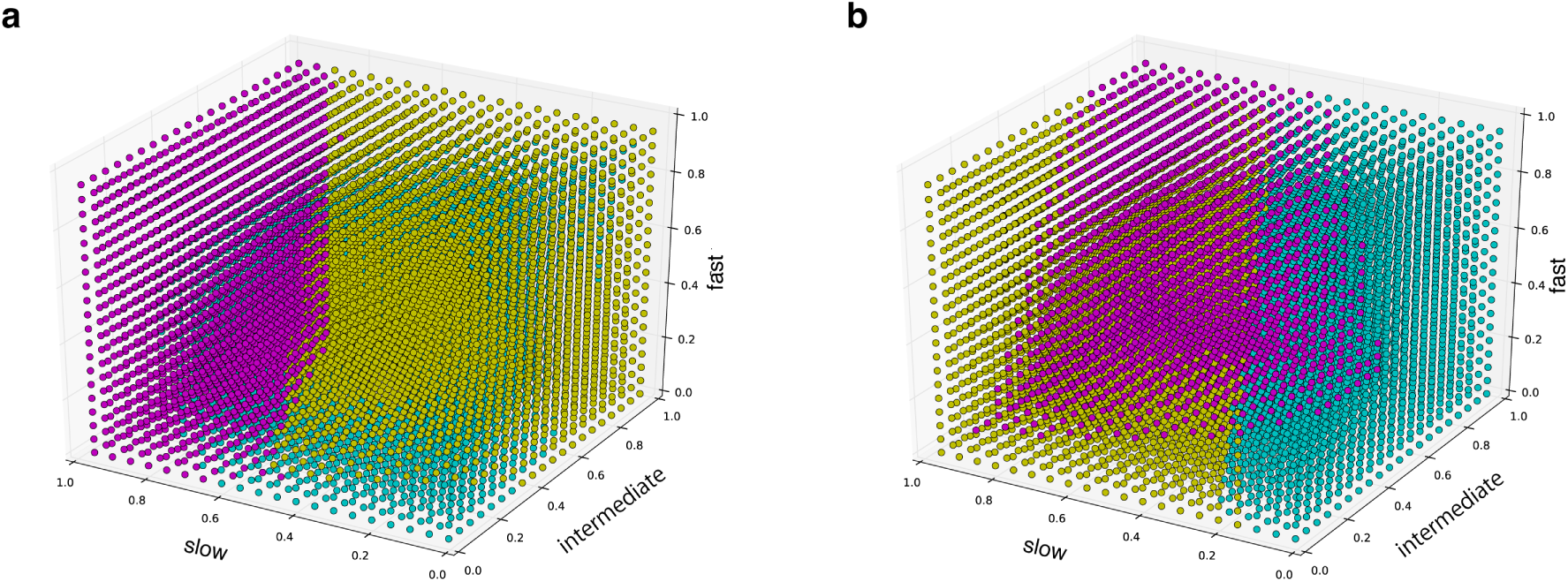
Initial conditions and the evolutionary fates in the infinite population size model. The relationship between the initial conditions and the stable equilibria the dynamics converge to in the three species community is illustrated when (a) *k* = 0.5 (small *k*) and (b) *k* = 1.5 (large k). The colors of the dots show which stable equilibrium the dynamics converges to when the initial fractions of generous individuals in each species are given by the coordinates of the dots. (a) cyan: the slow and the intermediate are selfish while the fast is generous. yellow: the slow and the fast are selfish while the intermediate is generous. magenta: the intermediate and the fast are selfish while the slow is generous. (b) cyan: slow is selfish while the others are generous. yellow the intermediate is selfish while the others are generous. magenta: the fast is selfish while the others are generous.

**Figure 7:**
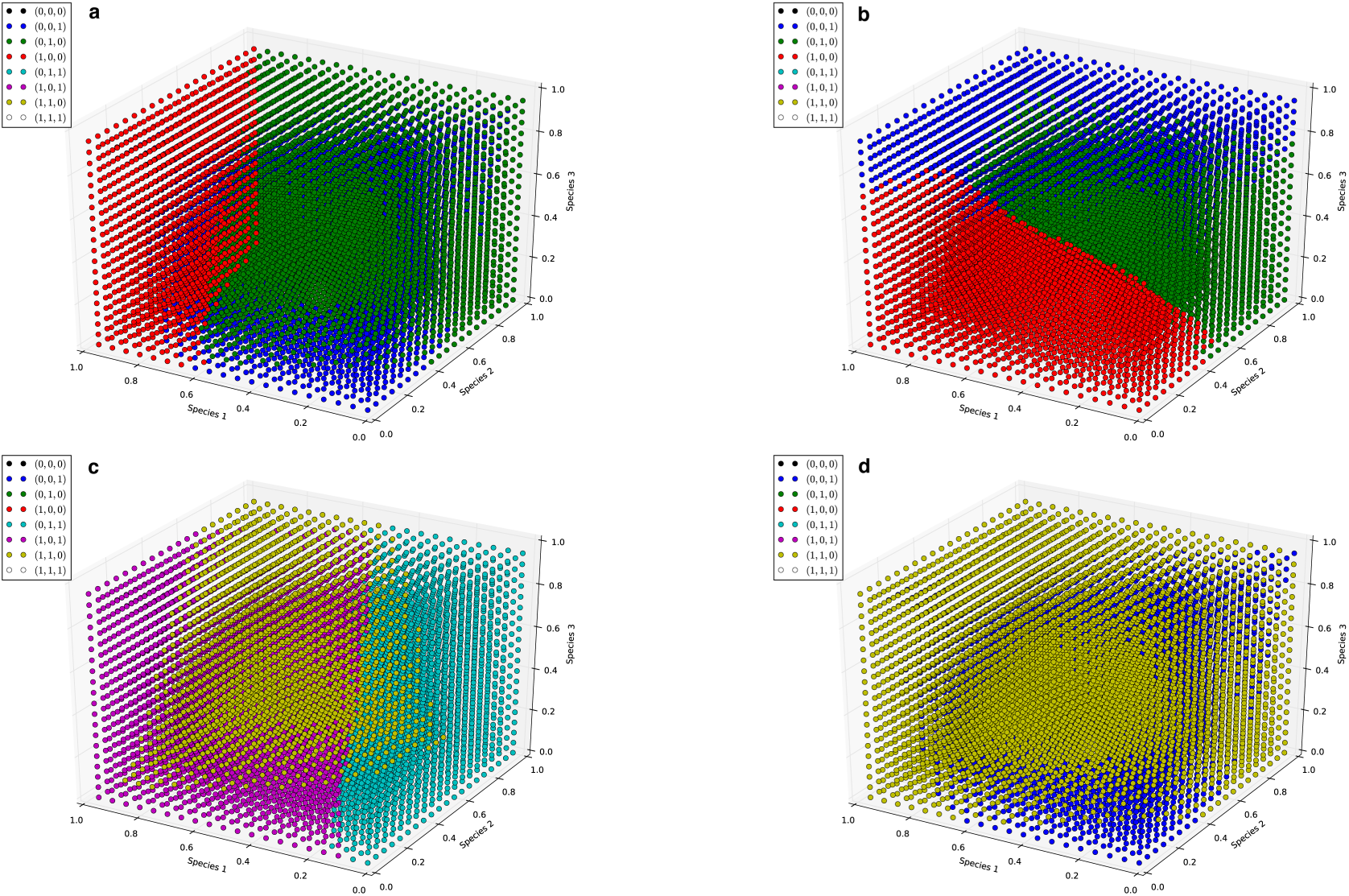
Examples of different population size model If *M* = 3, there exist 8 candidates of the stable equilibrium. Each dot indicates the initial fractions of generous individuals, and the color of the dots represents which equilibrium the dynamics converge to. While the evolutionary rate only changes the size of basin attraction, the value of *k* and the population sizes ***N*** can affect the stability of each equilibrium. In addition, the stable equilibria do not always the same number of generous species given the values of *k, M*, and ***N***. The parameters are (a): ***r*** = (1/8, 1, 8), *k* = 0.5, and ***N*** = (3, 3, 4), (b): ***r*** = (8, 1, 1/8), *k* = 0.5, and ***N*** = (3, 3, 4), (c): ***r*** = (1/8, 1, 8), *k* = 1.5, and ***N*** = (3, 3, 4), and (d): ***r*** = (1/8, 1, 8), *k* = 0.5, and ***N*** = (1, 1, 5)<ENT FONT=(normal text) VALUE=46>.</ENT>

